# Anaerobic oxidation of methane supports a minimal microbial community in a Subsurface Biofilm at Ginsburg Mud Volcano

**DOI:** 10.64898/2026.01.09.698606

**Authors:** Cleopatra Collado, Pedro Romero-Tena, Gunter Wegener, Marcus Elvert, Walter Menapace, Rafael Laso-Pérez

**Author notes:** These authors contribute equally to the work (Cleopatra Collado, Pedro Romero-Tena). Facultad de Veterinaria, Universidad Complutense de Madrid (UCM), Madrid, Spain. Ecologie Société Evolution, Université Paris-Saclay AgroParisTech, CNRS, Gif-sur-Yvette, France. **Correspondence:** Rafael Laso Pérez.

## Abstract

Deep marine sediments generate large amounts of methane, but most of this gas is consumed by the anaerobic oxidation of methane (AOM) mediated by microscopic consortia of anaerobic methane-oxidizing archaea (ANME) and sulfate-reducing bacteria (SRB). In this study, we investigated the AOM within a sulfate-methane transition zone (SMTZ) at a depth of ∼9.6 m at the rim of the Ginsburg mud volcano in the Gulf of Cádiz. The SMTZ is supplied with sulfate from both, overlying seawater and an underlying evaporitic deposit, and it coincides with a fracture zone that hosts a visible biofilm. Here, carbon dioxide shows the strongest ^13^C-depletion, indicating intense methane consumption. Metagenomic and lipid biomarker analysis of the biofilm revealed an exceptionally simple microbial community dominated by ANME-1b archaea (63%), which predominantly produce strongly ^13^C-depleted glycerol dialkyl glycerol tetraethers and, to a lesser extent, the less common macrocyclic archaeols. The putative partner bacterium Seep-SRB1c (*Desulfobacterota*) is less abundant (9%). Additionally, the biofilm contained five low-abundance heterotrophs that likely rely on biomass or metabolites released from the ANME-SRB consortium. Our study highlights the presence of active methanotrophic biofilms in subsurface sediments and suggests that these communities may play an overlooked role in mitigating seafloor methane emissions.

## Introduction

In marine sediments, microbial and thermogenic degradation of organic matter under anoxic conditions results in the production of large amounts of methane(1,2). However, most of this methane is oxidized in the sulfate methane transition zone (SMTZ), a process known as anaerobic oxidation of methane (AOM)(3). As a result, only small fractions of methane generated in the seabed reaches the water column, and eventually the atmosphere(4). AOM is typically mediated by syntrophic consortia of anaerobic methanotrophic archaea (ANME) and sulfate-reducing bacteria (SRB)(3). In this partnership, ANME completely oxidize methane to inorganic carbon via the reverse methanogenesis pathway, while SRB use the reducing equivalents derived from methane oxidation to reduce sulfate to sulfide(5,6). The syntrophic interaction between both partners seem to rely on direct electron transfer mediated by conductive structures, including extracellular cytochromes(7–9).

In recent years, the phylogeny and diversity of both partners has been studied in detail. ANME archaea are a polyphyletic group divided in three different clades (ANME-1, ANME-2, and ANME-3) closely related to methanogenic archaea of the *Halobacteriota*(5). Phylogenomic studies support ANME-1, also known as “*Candidatus* Methanophagales,” as an order-level group, while ANME-2 and ANME-3 are included within the family *Methanosarcinales*. Each group seem to inhabit different environments. ANME-1 can adapt to a wide temperature range, from psychrophilic to hyperthermophilic conditions(10). They are also highly abundant in chimney structures(11), and have been detected in terrestrial settings(5). Similarly, ANME-2 inhabits a wide variety of marine environments where concentrations of sulfate are slightly high, such as cold seeps, hydrothermal vents, and sediments, while ANME-3 predominantly inhabit submarine mud volcanoes and certain types of sporadic seepage(3,12).

SRB involved in the AOM include at least five clades of the phylum *Desulfobacterota* (HotSeep-1, Seep-SRB2, Seep-SRB1a, Seep-SRB1g and *Thermodesulfobacteria*)(6,10) that form syntrophic associations with ANME. These clades possess different features involved in the direct electron transfer from ANME to SRB such as multiheme cytochromes(6). Generally, Seep-SRB2 populations tend to associate with members of ANME-1 and ANME-2c(7), while Seep-SRB1a and Seep-SRB1g are exclusively related to ANME-2 clades(13,14). HotSeep-1 and *Thermodesulfobacteria* are found only in consortia with thermophilic ANME-1(10,15). These syntrophic SRB are phylogenetically related to other clades occurring in methane-rich sediments, but described as non-syntrophic, such as Seep-SRB1b, Seep-SRB1c, Seep-SRB1d, Seep-SRB1e, and Seep-SRB1f(6,16).

Within the often narrow SMTZs, AOM partners typically occur in small consortia invisible for the naked eye. Under high flux rates, AOM activity in surface sediments can lead to cell densities of 10^10^ ANME-SRB cells cm^-3^ (17). In the anoxic Black Sea, ANME-1 and ANME-2 were found to form dense microbial mats adhered to carbonate chimneys(18–20). Recent studies have identified AOM biofilms in subsurface sediments containing gas hydrates off the coast of Oregon, in the Indian Ocean, and in the Arctic Ocean(21–24). In general, these biofilms presented low diversity with predominance of ANME-1 archaea and SRB. In this study, we analyzed two biofilms embedded in a sediment core recovered from the foothill of the Ginsburg Mud Volcano (GMV), in the Gulf of Cadiz. The combination of metagenomics analysis and lipid and isotope geochemistry revealed a dense biofilm highly enriched on a single species of ANME-1b, that sustains a low-diversity microbial community.

## Study area

The Gulf of Cadiz (GoC), offshore the southwest Iberian Peninsula and northwest Morocco, is a geo-logically intricate and tectonically active region marking an oblique convergent plate boundary be-tween the African and Eurasian plates(25–27). It has become a key area for scientific research due to its widespread submarine fluid venting structures, particularly mud volcanoes (MVs) and pockmarks, which serve as pathways for fluids and sediments from several kilometers deep into the crust to the seafloor(28,29). These structures are significant for their geological, geochemical, and biological activity, their role in the Earth’s carbon cycle, and their potential as indicators for deep-seated pro-cesses, hydrocarbon exploration and geohazards(29,30).

Mud volcanism is prevalent in the GoC, with over 90 MVs identified(31,32). Their formation is closely associated with overpressurized methane and other fluids rising from deep source rocks and hydrocarbon deposits(28), alongside halokinetic movements and a compressional tectonic regime that mobilizes sediments to the surface(33,34). Hydrocarbon gases from GoC MVs typically have a deep thermal origin, often from Mesozoic (Jurassic and Cretaceous) source rocks(35,36), displaying biogenic components and complex subsurface migration patterns(35,37–39). Fluid migration path-ways are typically fault-controlled, utilizing various fault types to reach the surface (40,41).

The GMV is among the largest mud volcanoes in the GoC, rising 280 m above the seafloor and spanning up to 3,860 m in diameter (31,32,42). Its complex architecture, with stacked edifices and intrusive complexes, results from alternating mud extrusion events and dormancy periods, being par-ticularly active since the Late Pliocene(43). A recent study revealed the complex fluid pathways within the GMV, distinguishing between the provenance of summit and moat. Summit fluids primari-ly originate from clay dehydration within the AUGC, and are channeled by central conduits with high advection velocities, while moat fluids have slower advection velocities and additional geochemical effects from evaporite dissolution (44). Peripheral seepage at MV edifices is linked to fractures formed by edifice subsidence(44). While current overall methane emission rates from GoC MVs are generally low, the sustained advective fluid transport and complex fluid circulation at Ginsburg im-ply continuous, albeit variable, methane expulsion, making it an important site for research into deep-sourced carbon cycling (39,40,46). The samples analyzed in this study come from the sediment core GeoB23047-3 at the foothills of GMV, and associated with peripheral seepage.

## Material and methods

### Sample collection and geochemistry

Sediment samples analyzed in this study were collected during the R/V METEOR Cruise M149 in 2018(45) in close proximity to the Ginsburg Mud Volcano (35°22.863’ N, 7°04.128’ W). The GeoB23047-3 sediment core, of 40.30 meters length, was obtained for geological analysis using the seafloor drill rig MeBo at 1126 m water depth(45). A fractured crosscut of the core was located at 9.45-9.70 m depth which was filled with a slimy dark-brown organic biofilm. Four samples, including biofilm samples 1 (9.50-9.55 m) and 2 (9.65-9.68 m) as well as sediments samples 1 (9.44-9.47 m) and 2 (9.60-9.63 m), were collected. These samples were processed on the ship and stored under anoxic conditions at −20°C for analysis.

Pore water sampling was carried out on sediment core whole-round sections onboard, with samples taken every 40 cm with Rhizon samplers(45) (5 cm length, 0.15 μm porous polymer; Rhizosphere Research Products), soaked in distilled water before use. Small holes were drilled in the core liners to insert the Rhizons into the sediments. A vacuum was applied to the Rhizons using medical syringes(46). Pore water samples for anion concentrations measurements were stored in 2 mL air-tight Eppendorf® vials at 4 °C until their post-cruise analyses at MARUM. Pore water SO ^2−^ concentrations were determined by ion chromatography (Metrohm 861 Advanced Compact IC, Metrohm A Supp 5 column, 0.8 mL min−1, conductivity detection after chemical suppression) in samples diluted 1:40 with Milli-Q-grade H_2_O. The detection limit for SO_4_ ^2−^ was 0.5 µM, with a precision of <1%.

Sediment samples for gas analysis were taken by removing approximately 3 cm³ of sediment from the lower part of each core section. Each sediment plug was transferred to a glass container containing 10 ml of a 1 M KCl solution and sealed with a rubber stopper and a metal sleeve. At MARUM, the δ^13^C values of methane and CO_2_ in the headspace were determined using a Trace GC connected to a DELTA Plus IRMS via a GC combustion interface III (all Thermo Finnigan, Bremen, Germany) as described previously(47). In brief, the GC was equipped with a Supelco Carboxen 1006 Plot capillary column (30 m × 0.32 mm i.d.) and He as carrier gas at a constant flow rate of 2 ml min^-1^. The injection was performed in split mode, and the oven temperature of 40 °C (held for 2 min) was increased at a rate of 40°C per min to 240°C, where it was maintained for 3 min. δ^13^C values are reported in the delta notation relative to the Vienna Pee Dee Belemnite (VPDP) standard. Primary standardization of the carbon isotope analyses was based on multiple injections of in-house reference CO_2_ with a known δ^13^C value at the beginning and the end of each analytical run. Standard deviation of repeated analyses of the samples was <1.0‰.

### DNA extraction, preparation of 16S rRNA gene libraries, and tag sequencing

DNA was extracted from the samples with a DNeasy Power Water kit (Qiagen, Germany). Amplicon libraries were prepared by following the 16S metagenomic sequencing library preparation guide provided by Illumina with primers targeting the V3-V4 region for bacteria (Bact0341-Bact0785) and V3-V5 for archaea (Arch349F-Arch915R). The gene amplicon libraries were sequencing using Illumina MiSeq 300 platform (Illumina, U.S.A.) at the Center for Biotechnology (CeBiTec, Bielefeld, Germany). The retrieved sequenced were processed as previously described (See Supplementary Information)(48).

### Metagenomic analysis and metabolic prediction

The extracted DNA was also used for metagenomic sequencing using Illumina MiSeq 300 machines at CeBiTec. The metagenomic reads were trimmed using BBduk (qtrim=t; trimq=20; minlength=150)(49), and then coassembled using SPAdes (v3.15.5) with default parameters(50). Assembled contigs were automatically binned using Maxbin2 v2.2.7(51) with default parameter and Metabat v2.2.15(52), selecting archaeal and bacterial-specific marker genes (markerset = 40). The results of the automatic binning were used as a guide for a manual binning process using Anvi’o v.8.0(53). The resulting bins were refined via targeted reassembly following an approach previously described(54) and the best reassembly was selected as Metagenome-Assembled Genomes (MAGs). The quality, taxonomy and metagenomic coverage of the MAGs were respectively evaluated with CheckM v1.1.3(55), GTDB-Tk v2.1.1(56) and CoverM v0.7.0(57).

Open reading frames (ORFs) were predicted and annotated using Prokka(58). Protein annotation was also performed using DIAMOND v.2.1.9.163(59) against the COGs(60), ArCOGs(61) and KEGG(62), and the MEROPS(63) databases. Moreover, HMMER(64) was used against the Pfam(65) and TIGRfam(66) databases. Multiheme cytochrome were determined by counting the number of “CXXCH” motifs. Hydrogenases were identified using the HydDB hydrogenase database(67).

### Phylogenetic and phylogenomic analysis

To determine the phylogenomic position of ANME-1-GMV a phylogenomic tree was constructed using 105 MAGs from the class *Syntropharchaeia* (Table S1). Using Anvi’o v8.0, a protein alignment of 31 single-copy gene markers (Table S2) was generated with the command: ‘anvi-get-sequences-for-hmm-hits --return-best-hit --max-num-genes-missing-from-bin 7’. The concatenated file was used to calculate the phylogenomic tree with RAxML v8.2.13(68) adding the corresponding partition file and following parameters: ‘raxmlHPC-PTHREADS -m PROTGAMMAAUTO -f a -N autoMRE -p 45670 -k -x 5789’. The tree was visualized and modified with iTOL (https://itol.embl.de/). Besides, we compared the average nucleotide identities (ANI) of a selection of genomes phylogenetically close to ANME-1-GMV using JSpeciesWS (https://jspecies.ribohost.com/jspeciesws/).

To determine the evolutionary position of the ANME-1-GMV McrA, a phylogenetic tree was constructed using an alignment obtained with MAFFT v7.525(69). The phylogenetic tree was inferred using RAxML(68) v8.2.13 with the parameters ‘raxmlHPC-PTHREADS -m PROTGAMMAAUTO -f a -N 1000 -p 253615 -k -x 45272’. Tree visualization was performed using ITOL. Additionally, a synteny analysis was made around the genes encoding for the MCR enzyme for selected genomes (Table S1) using CAGECAT program(70).

For studying the evolution of GDGT-synthases we used the GrsA (SACI_RS01165) and GrsB (SACI_RS07560) GDGT-synthases protein sequences, as query sequences from *Sulfolobus acidocaldaricus* in BLAST to look for homologs (e-value < 1e-20, identity >20%, number of hits=100) in the 105 MAGs of the class *Syntropharchaeia*. Redundant sequences (>90% similarity) were removed using CD-HIT v4.8.1.(71). Then, we introduced our sequences in an alignment of GDGT-synthases like proteins from a previous study(72) (Table S3) using MUSCLE v5.1(73) The phylogenetic tree was calculated using RAxML as previously described.

### Lipid analysis

Two samples (Biofilm 1, Sediment 1) were investigated for the distribution and carbon isotope composition (δ^13^C values) of lipid biomarkers. We targeted the full suite of intact polar lipids (IPLs) and core lipids as well as polar-lipid derived fatty acids (PLFAs) and neutral lipids, including hydrocarbons and alcohols. Briefly, a total lipid extract (TLE) was obtained according to Sturt et al. (2004)(74) based on a modified Bligh & Dyer protocol. Before extraction, 1 μg each of 1,2-diheneicosanoyl-sn-glycero-3-phosphocholine, 5α-cholestane, *n*-nonadecanol and 2-methyloctadecanoic acid was added as internal standard. In a TLE aliquot, the PLFAs were converted to fatty acid methyl esters (FAMEs) by saponification with KOH/MeOH followed by derivatization with BF_3_/MeOH(75). Neutral lipids obtained during the saponification reaction were derivatized with bis(trimethylsilyl)trifluoroacetamide prior to analysis, yielding TMS-derivatives(76). IPL and core lipid fractions were separated from an aliquot of the TLE using preparative liquid chromatography(77) and analyzed separately (see below). Ether lipids in the separated IPL and core lipid fractions were subjected to ether cleavage using BBr_3_ followed by reduction with lithium triethylborohydride, forming hydrocarbons(78).

Neutral lipids, FAMEs and ether-cleaved hydrocarbons of the core and IPL fractions were analyzed by gas chromatography–flame ionization detection for quantification (GC-FID; Thermo Finnigan TRACE GC) and GC-mass spectrometry for identification (GC-MS; Thermo Finnigan TRACE GC coupled to a TRACE MS) using columns and temperature program settings as previously described(79). Similarly, δ^13^C analysis was performed using GC-isotope ratio MS (GC-IRMS; Thermo Scientific TRACE GC coupled via a GC IsoLink interface to a DELTA V Plus). δ^13^C values are reported in the delta notation relative to VPDP and are referenced to an in-house laboratory CO_2_ gas with an analytical precision better than 1‰ as determined by long-term measurements of an *n*-alkane standard with known isotopic composition of each compound. δ^13^C values of FAMEs and TMS-derivatives were corrected for carbon introduced during derivatization.

IPL and core lipid fractions were analyzed by coupled ultra-high performance liquid chroma-tography - mass spectrometry (UHPLC-MS; Dionex Ultimate 3000 UHPLC instrument coupled to a Bruker MaXis ultra-high resolution quadrupole time-of-flight mass spectrometer). IPLs were subjected to hydrophilic interaction LC coupled with electrospray ionization-MS(80) and core lipids were analyzed by two coupled Waters Acquity UPLC BEH Amide columns and atmospheric pressure chemical ionization-MS(81). For the core lipid analysis, an injection standard (1 ng of C_46_-GTGT) was added to the sample and compounds were quantified based on response factors of the detected core structures.

## Results and discussion

### Sampling and geochemistry

The sediment core consisted mostly of light grey foraminifera-rich nannofossil ooze, consistent with a hemipelagic origin rather than mud-volcano material. Between 9.45 m and 9.70 m depth, the core presents a crack (Figure 1A), where SO_4_ ^2−^ concentrations approach 0 mM (Figure 1B) and pyrite concretions and precipitates are present(45), marking the SMTZ. Above and below this interval, sulfate concentrations increased, indicating sulfate supply from the overlying water column as well as upward, deep-rooted sulfate flux from underlying evaporitic sediments. The crack was filled with a slimy dark-brown organic macroscopic biofilm (Figure 1A). Along the core, methane headspace gas δ^13^C values were typically about −65‰, but decreased to −78‰ in the sulfate depleted interval (Figure 1B). Similarly, CO_2_ δ^13^C values declined from −25‰ to as low as −40‰. At this depth, Xu et al. (2025) identified strong seismic reflectors with reverse polarity, suggesting active fluid seepage(44).

**Figure 1.**
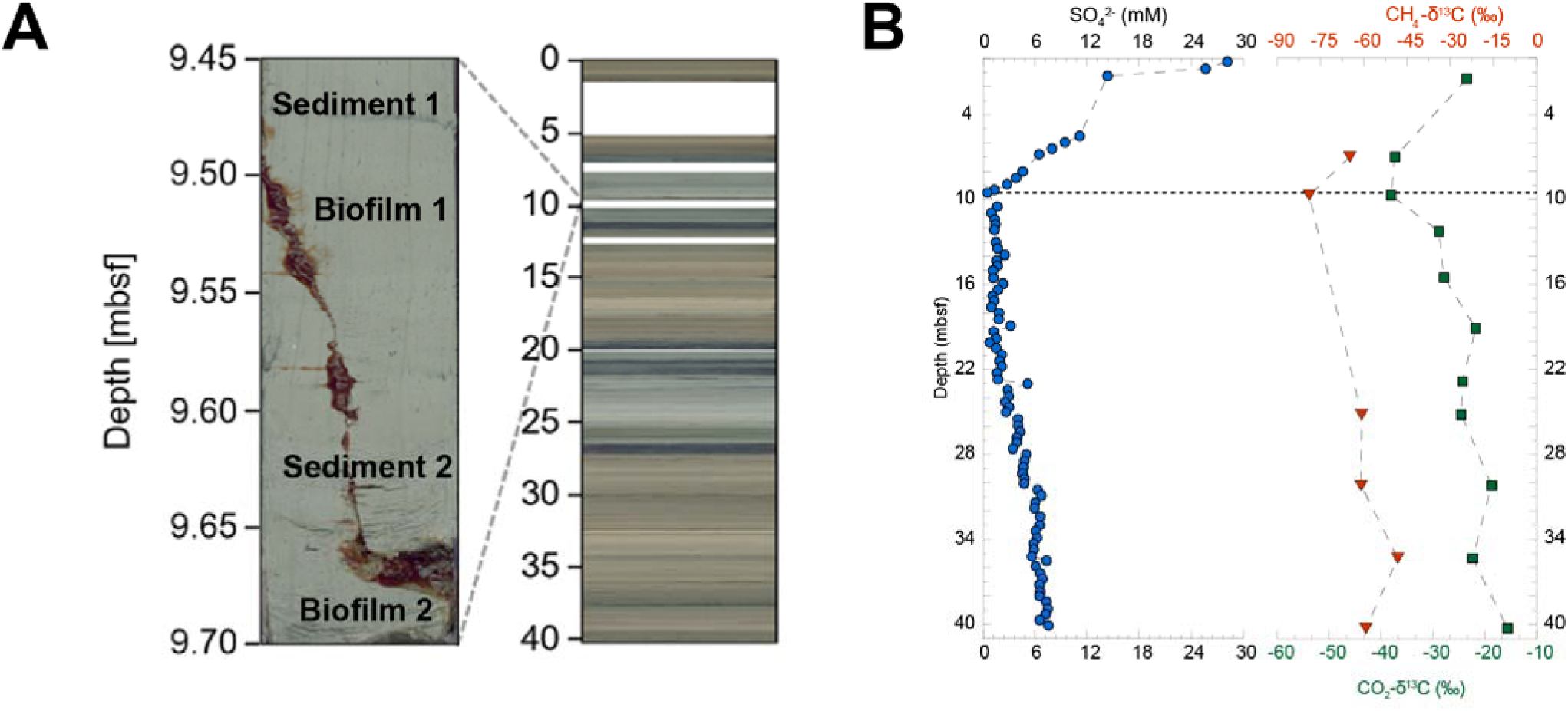
Sampling and geochemical information. (a) Photography of the GeoB23047-3 sediment core and the corresponding biofilms. (b) Geochemical characterization of the GeoB23047-3 sediment core. Concentration of SO_4_ ^2−^ in mM (blue circles) and δ^13^C values of methane (red triangles) and carbon dioxide (green squares) over depth. The horizontal dotted line indicates the location of the biofilm samples.

### A few monoclonal taxa dominate the biofilm

To resolve the nature of the biofilm, we extracted the DNA and compared this with two nearby sediment samples (Figure 1A). Whereas the sediment samples contained very little DNA (∼0.6 ng per gram of sediment), we extracted up to 1.1-1.4 µg of DNA from the microbial biofilm. According to 16S rRNA gene amplicon sequencing, both biofilm samples presented an exceptionally low microbial diversity (Figure 2). Their archaeal 16S rRNA libraries were dominated by ANME-1 (>95%), with a small proportion of an Asgard archaeon (Figure 2A). The biofilm bacterial 16S rRNA gene libraries contained sequences of four families with similar relative abundances: *Bacteroidetes, Calditrichaceae, Desulfobacterota* and *Spirochaetaceae*. In the surrounding sediments, the ANME-1b represent about 40 to 60% of the archaea, while the bacterial community was much more diverse and very different compared to the biofilm community with groups like *Gammaproteobacteria, Bacilli* and *Actinobacteria* dominating the sample.

**Figure 2.**
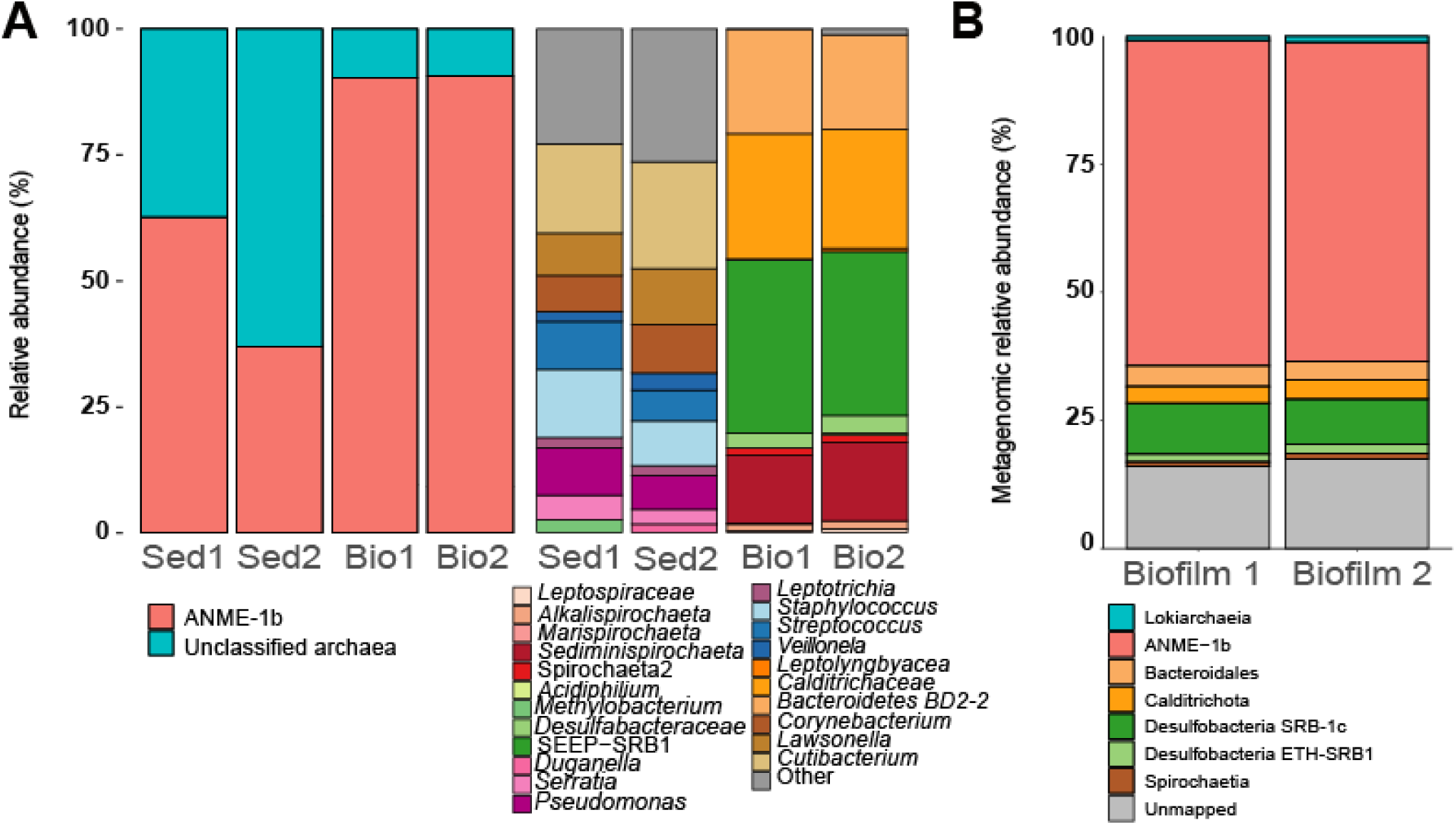
Microbial community structure of the biofilms. (A) archaeal (left) and bacterial (right) 16S rRNA gene-based profiles of the biofilm and sediment community. (B) relative metagenomic abundance based on read mapping to metagenome-assembled genomes (MAGs). ANME-1 dominates the community (63.5% in Biofilm 1 and 62.3% in Biofilm 2) followed by Seep-SRB1c. Sed: Sediment. Bio: Biofilm

To analyze the function of the biofilm community, we performed metagenomic sequencing of the biofilm samples. We retrieved seven metagenomic-assembled genomes (MAGs) that represent all taxa observed in the amplicon libraries of the biofilms (Table 1, Figure 2B). Two MAGs correspond to archaea, one affiliated to the ANME-1 (ANME-1-GMV) and recruiting up to 60% of the metagenomics reads, and the second closely related to the recently described *Prometeoarchaeum syntrophicum* (class *Lokiarchaeia*)(82). The five bacterial MAGs include two members of the class *Desulfobacteria* (Seep-SRB1c-GMV with 9% of the reads and ETH-SRB1-GMV with less than 2% of the reads). Other bacterial genomes were affiliated to the classes *Spirochaetia, Calditrichia* and *Bacteroidia* with relative abundances between ≤1% and 4%.

**Table 1.** Summary of metagenome-assembled genomes (MAGs) recovered from the biofilm samples. Metrics include genome completeness and contamination (as percentage), N50 (as pb), genome size (pb), proportion of metagenomic reads mapping to the MAG and the corresponding taxonomy according to GDTB. The proportion of unmapped reads for each sample was 16.07% for Biofilm and 17.52% for Biofilm 2. Comp: Completeness. Cont: Contamination. MR: Mapped reads

| MAG name | Co mp (%) | Co nt (%) | N50 (pb) | Geno me Size (pb) | MR Biofil m 1 (%) | MR Biofil m 2 (%) | GTDB Classification |
| --- | --- | --- | --- | --- | --- | --- | --- |
| ANME-1-GMV | 91.45 | 1.25 | 42867 | 2063818 | 63.54 | 62.29 | d__Archaea;p__Halobacteriota;c__Syntropharchaeia;o__ANME-1;f__ANME-1;g__JAFNKG01;s__ |
| Bacteroidales-GMV | 96.77 | 1.52 | 92771 | 4114521 | 4.01 | 3.69 | d__Bacteria;p__Bacteroidota;c__Bacteroidia;o__Bacteroidales;f__VadinHA17;g__JADFUQ01;s__ |
| Calditrichia-GMV | 84.66 | 0.4 | 135459 | 4015801 | 3.29 | 3.65 | d__Bacteria;p__Calditrichota;c__Calditrichia;o__f__g__s__ |
| Seep-SRB1c-GMV | 96.05 | 2.0 | 181442 | 3961353 | 9.92 | 8.81 | d__Bacteria;p__Desulfobacterota;c__Desulfobacteria;o__C00003060;f__C00003060;g__C00003060;s__ |
| ETH-SRB1-GMV | 70.26 | 2.63 | 38099 | 3028958 | 1.54 | 1.82 | d__Bacteria;p__Desulfobacterota;c__Desulfobacteria;o__Desulfobacterales;f__ETH-SRB1;g__s__ |
| Loki-GMV | 86.09 | 1.85 | 46426 | 3644098 | 0.82 | 1.17 | d__Archaea;p__Asgardarchaeota;c__Lokiarchaeia;o__CR-4;f__MK-D1;g__s__ |
| Spirochaetia-GMV | 85.3 | 1.1 | 28697 | 3533252 | 0.80 | 1.05 | d__Bacteria;p__Spirochaetota;c__Spirochaetia;o__DSM-16054;f__Sediminispirochaetaceae;g__s__ |

### ANME archaeal lipids dominate the biofilm

We further investigated the lipid biomarker composition and corresponding δ^13^C values in one of the biofilm samples (biofilm 1) in comparison to one of the surrounding sediments (sediment 1). In the surrounding sediments, core GDGTs reached concentrations up to 11 µg g_dw_ ^−1^ for GDGT-0 (Table S4) and DAGEs had values between 0.3 and 5.4 µg g_dw_ ^−1^. There were only traces for archaeal IPL (PG-GDGT-0: 0.033 µg g_dw_^−1^), whereas bacterial IPLs were undetectable (Table S4). This dominance of core lipids over intact compounds is typical for deep-sea sediments, which often contain larger amounts of fossil lipids, but much less IPLs as marker for alive cells(83). In contrast, the biofilm contained much higher concentrations across all lipid classes, dominated by intact and core glycerol dialkyl glycerol tetraethers (GDGTs). These accounted for 88% of total IPLs (PG-GDGTs) and 55% of total core lipids (GDGTs), with individual GDGTs reaching concentration of ∼1,5 mg g ^−1^ (Figure 3A-B; Table S4). The most abundant compounds were GDGTs without or with one cyclopentane ring (i.e. PG-GDGT-0, GDGT-0 and GDGT-1), strongly suggesting that the biofilm was dominated by ANME-1 archaea(84). Besides, we detected substantial amounts of intact and core lipids of archaeol, as well as its unsaturated and macrocyclic counterparts. Archaeol was most abundant with up to 298 µg g_dw_^−1^. From a biosynthetic perspective, phosphatidyl archaeols with a PG headgroup are considered the most likely precursors of ANME-derived PG-GDGTs whereas intact macrocyclic PG-archaeol is assumed to be a side product of the tetraether synthase (Tes) during GDGT formation(85,86). We also observed a large suite of unsaturated and macrocyclic PE archaeols with up to four double bonds and potentially three cyclopentane rings. However, we were unable to detect any corresponding PE-GDGT, suggesting that PE-archaeols probably do not serve as precursors for PE-GDGTs, but rather have another function in the cytoplasmic membrane.

**Figure 3.**
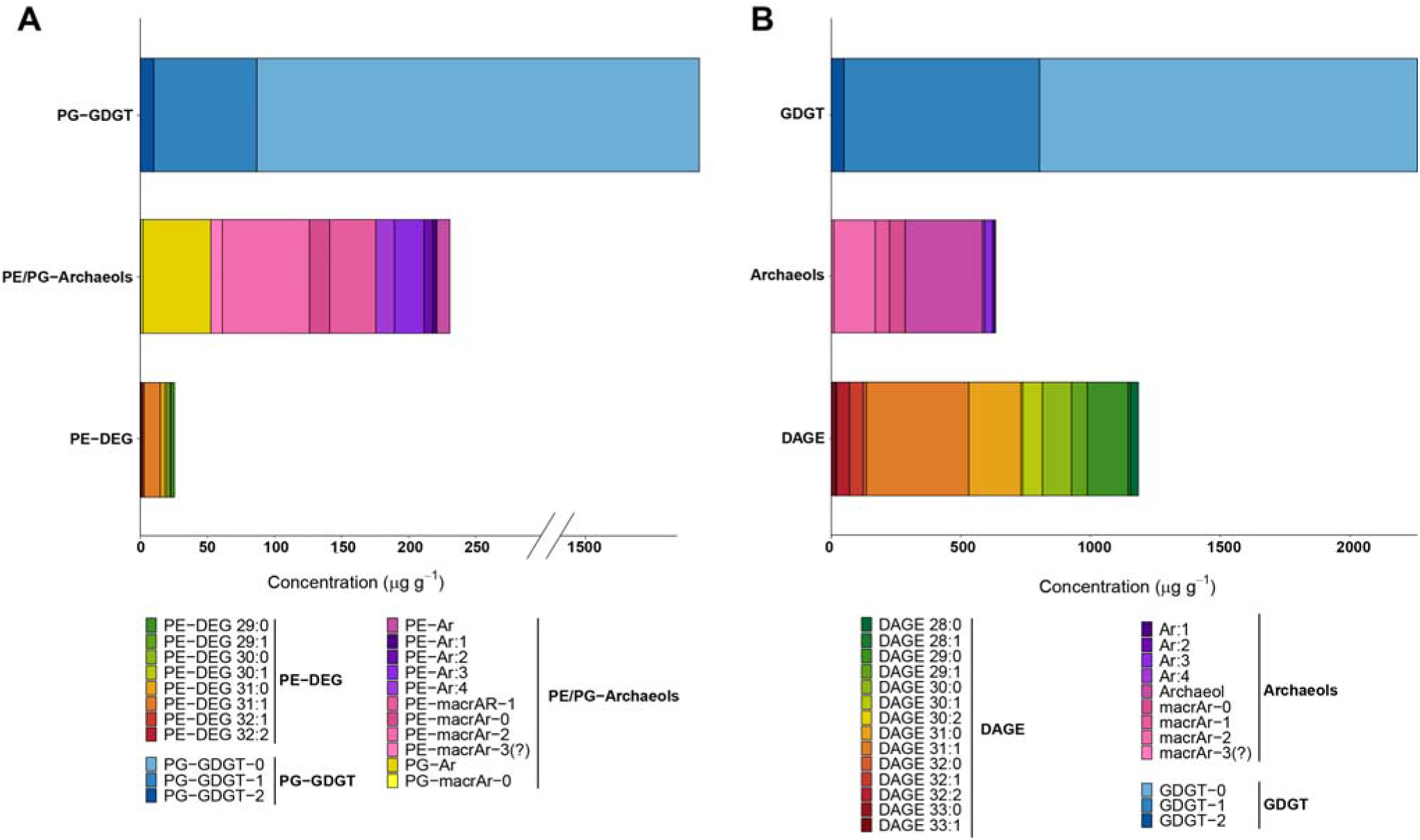
Concentrations of (A) intact polar lipids (IPLs) and (B) core ether lipids from the Biofilm 1 sample. ANME-1 characteristic lipids are dominating (PG-GDGTs: 88% of total IPLs, GDGTs: 55% of total core lipids) with up to ∼1.5 mg g_dw_^−1^ found for PG-GDGT-0 and GDGT-0. Bacterial-derived diether lipids are more abundant in the core lipid fraction, making up to 29% of the total core lipids. PE-DEG: phosphatidylethanolamine diether glycerol, PE-Ar: phosphatidylethanolamine archaeol, PE-macrAr: phosphatidylethanolamine macrocyclic archaeol, PG-Ar: phosphatidylglycerol archaeol, PG-macrAr: phosphatidylglycerol macrocyclic archaeol, PG-GDGT: phosphatidylglycerol glycerol dialkyl glycerol tetraether, DAGE: dialkyl glycerol ether.

The δ^13^C analyses of free archaeols and of the ether cleavage products biphytane and phytane, both obtained from intact and core GDGTs and archaeols, yielded δ^13^C values as low as −102‰ (mean: −98.8‰; Table 2, Table S5), corroborating active AOM mediated by ANME-1(87–89). We did not detect any of the characteristic isoprenoidal hydrocarbons that occur in many other AOM systems, such as crocetane or pentamethylicosane, but squalene with a δ^13^C value of −97‰ (Table 2). Although squalene is not commonly reported as a diagnostic lipid of ANME communities(88,90), its very low δ^13^C value here is in line with all of the other archaeal lipids ascribed to ANME-1. In the sediment sample, archaeal lipids exhibited much more positive δ^13^C values, indicating the presence of additional archaeal linages such as the *Lokiarchaeia* detected in this study (Figure 2, Table 1), whose lipids likely mix with those of ANME-1, and thereby dilute the strong AOM isotopic signal. Indeed, biphytane chains of GDGTs of *Prometheoarchaeum*, from the same genus detected here, contain Bp-0 to Bp-2 moieties(91). We also detected a signal from remnants of planktonic archaea in the deep MV sediment, namely crenarchaeol, with a δ^13^C value of ∼−15‰ measured for the ether cleavage product Bp-cren (Table 2, Table S5).

**Table 2.**
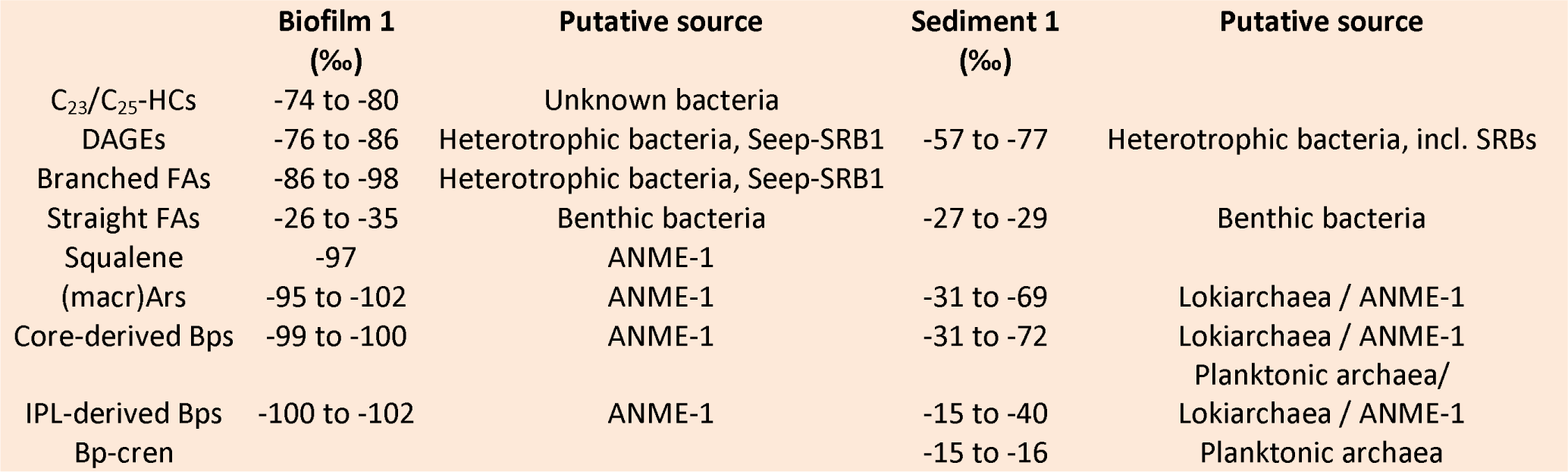
Overview of δ^13^C value ranges of lipid biomarker groups detected in the Biofilm 1 and Sediment 1 sample and their putative sources. Detailed information on each lipid biomarker accessible for δ^13^C analysis can be found in Table S5. HCs: hydrocarbons, DAGEs: dialkyl glycerol ethers, FAs: fatty acids, (macr)Ars: (macrocyclic) archaeols, Bps: biphytanes with 0, 1 or 2 cyclopentane rings, Bp-cren: biphytane with 2 cyclopentane and 1 cyclohexane ring derived from crenarchaeol.

The only bacterial IPLs were PE-DEGs with a total concentration of ∼26 µg g_dw_ ^−1^ (Figure 3A). The PE-DEGs contained single alkyl side chains with 14 to 16 carbon atoms (Table S4). These IPLs might be produced by the partner bacterium Seep-SRB1 or heterotrophic bacteria, and its lower concentration would imply a minor abundance of active bacteria (Table 1). In contrast, the core lipid fraction contains more bacterial lipids, in particular the dialkyl glycerol ethers (DAGEs) 31:1, 30:1, 30:0 and 29:0 with concentrations of up to ∼400 µg/g dry weight (Figure 3B), representing 29% of the total core lipids. This exceeds the estimates for metagenome-based abundances for bacteria (Table 1). Like the PE-DEGs, the DAGEs featured alkyl side chains between 14 and 16 carbon atoms, and based on published information(92–95) are dominated by monunsaturations and anteiso-branching (Table S4). DAGEs have been repeatedly reported from AOM environments(92,93,95) and are generally interpreted to originate from heterotrophic bacteria, including SRBs(94,96). However, enrichment studies also indicate that autotrophic SRB partners in AOM consortia can produce a subset of these DAGEs(97,98). We did not detect any ester-based IPLs but were able to obtain fatty acid signals after TLE saponification, predominantly branched FAs, although in lower concentrations than the DAGEs. The δ^13^C values of the branched FAs and of the DAGEs, analyzed from the neutral lipids, were also strongly ^13^C-depleted, though slightly less negative than the archaeal lipids, with mean values of −89.9‰ and −80.6‰ respectively (Table 2, Table S5). These results indicate a substantial flux of methane-derived carbon into these bacterial compounds. Notably, the more negative ^13^C values among the fatty acids, especially the iso-branched fatty acids *i*C16:0 with a value of −98‰, approach those of archaeal lipids. This underscores the presence of heterotrophic bacteria in the microbial community (i.e., *Spirochaetia, Calditrichia,* and *Bacteroidia*; Figure 2, Table 1), which are not directly involved in AOM. Instead these heterotrophy likely feed on ^13^C-depleted necromass or cell exudates/lysates produced by the AOM core community, possibly involving branched-chain amino acids that serve as primers of branched FA biosynthesis(99).

The biofilm neutral lipids also contain substantial amounts of C_23_ as well as undescribed C_25_ hydrocarbons with up to 2 double bonds (in sum 380 µg g_dw_^−1^, Table S4). Such hydrocarbons have been reported before in AOM environments, especially when ANME-1 is enriched(88,92,100), and were putatively assigned to bacteria, although their function remains unknown. In the biofilm, these compounds display a third, more positive δ^13^C range of −74‰ to −80‰ of lipid biomarkers (mean value: −76.4‰, Table 2). Due to their absence in the surrounding sediment, we propose that these hydrocarbons represent biomarkers for bacteria that are specific to ANME-1-dominated AOM systems, particularly those where abundant archaeal biomass supports heterotrophic activities, such as in microbial mats(100).

### ANME-1-GMV genome host two *mcrA* genes

ANME-1-GMV MAG affiliated to the ANME-1b clade, which includes among others the genera “*Candidatus* Methanophaga” and “*Candidatus* Methanoalium”(5), although it does not belong to any of these genera. Instead, ANME-1-GMV has an ANI value over 97% with the previously described genome JAFNKG01 sp. 030602585 (GCA_030602585.1) from a global cold seep genome catalogue (Table S6)(101). Together, these MAGs form a distinct clade related to “*Candidatus* Methanophaga*”* (Figure 4A), a genus typically associated with cold seep environments(54).

**Figure 4.**
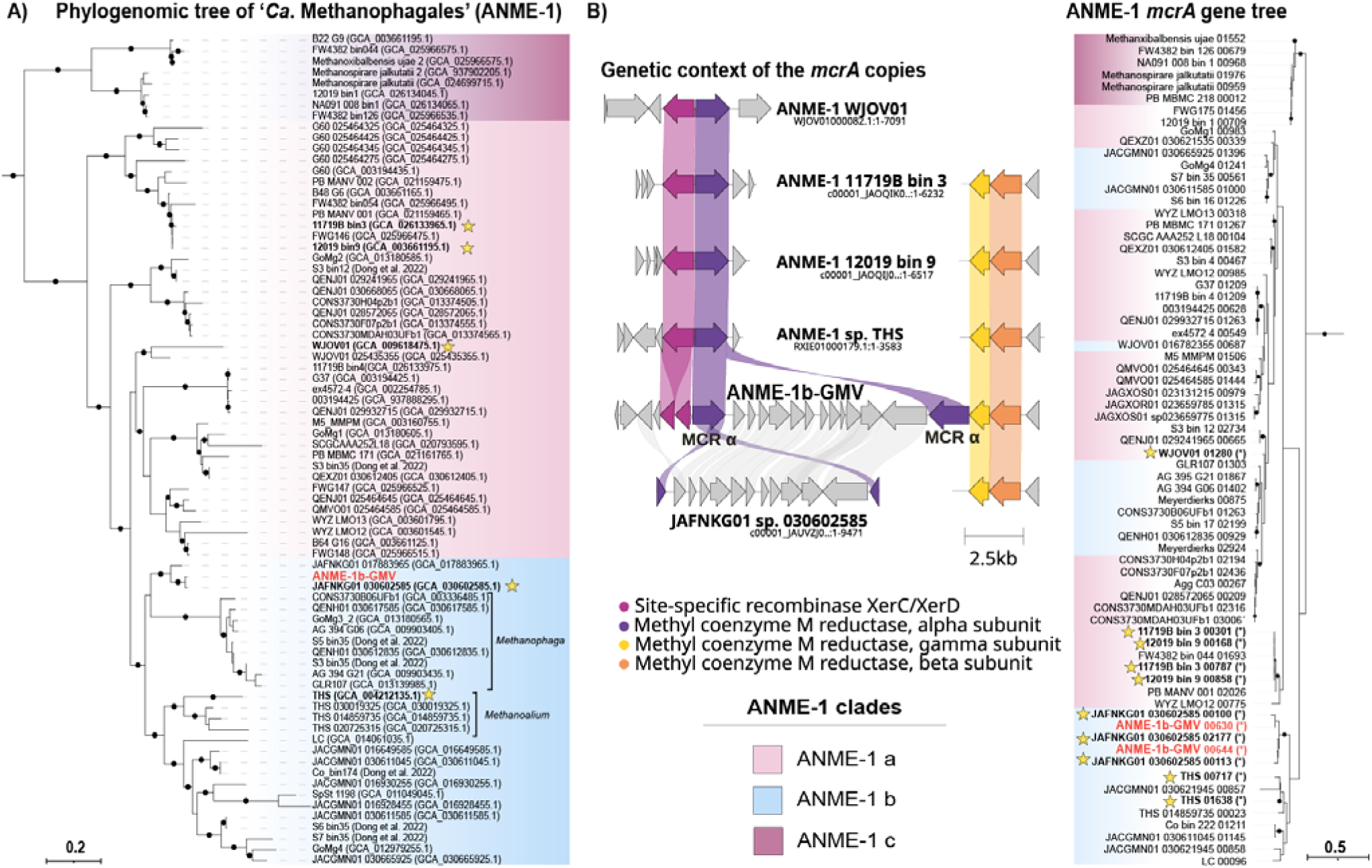
(A) Maximum-likelihood phylogenomic tree of the ANME-1 (“*Ca.* Methanophagales”). The outgroup (*Syntropharchaceae*) is not shown. (B) Phylogenetic tree of *mcrA* genes from ANME-1 genomes and genetic context of the *mcrA* gene copies of interest. The tree was constructed from aligned McrA amino acid sequences using maximum likelihood inference. Labels written in bold and stars indicate genomes with truncated *mcrA* genes, or the corresponding truncated genes. ANME-1b-GMV is marked in red. Filled circles indicate bootstrap values >90%.

**Figure 5.**
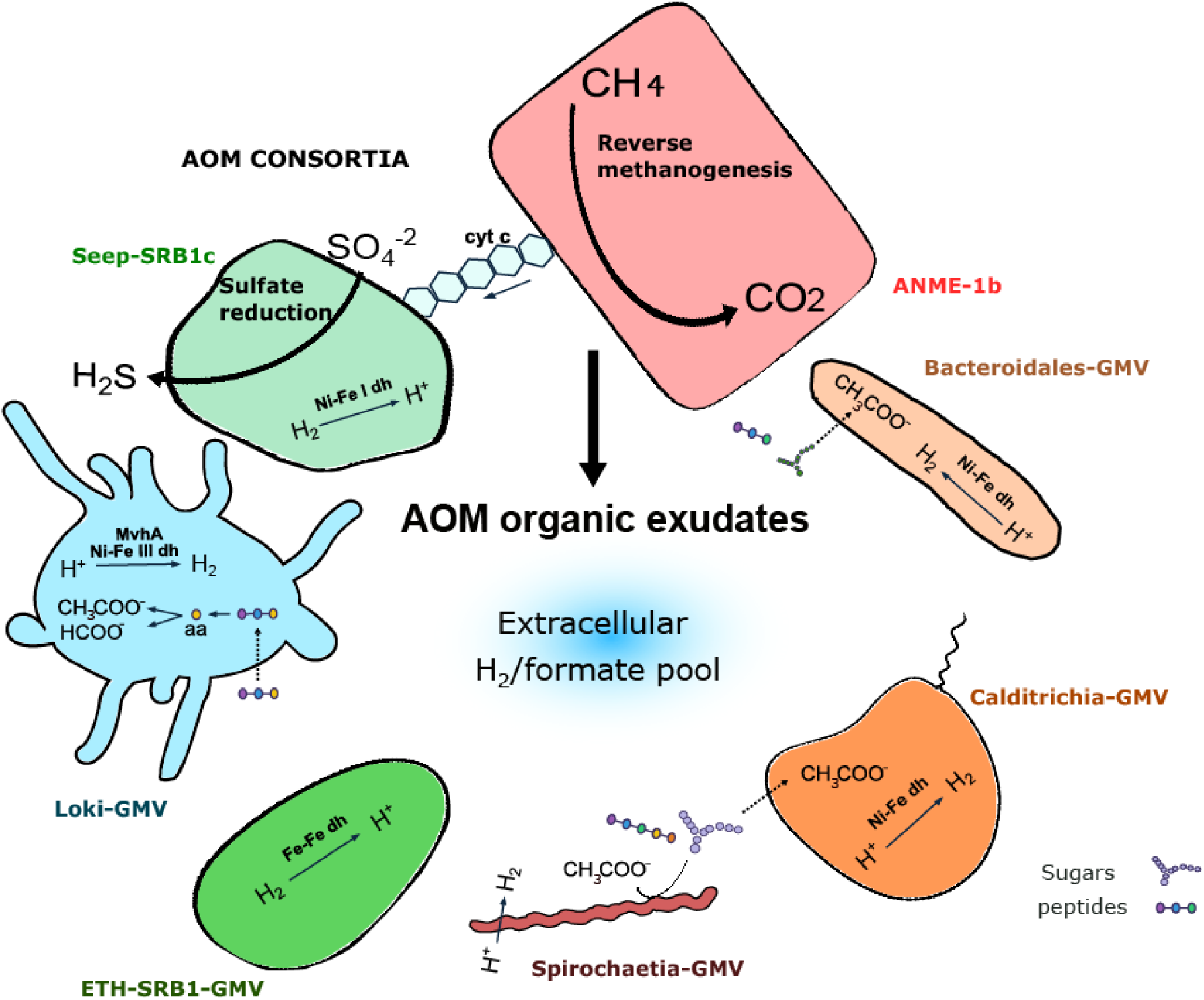
Overview of key functional traits in the biofilm MAGs in their environmental context. The model suggests that biomass or metabolites produced by the ANME and their partner bacteria (polysaccharides, peptides) serve as energy and carbon source for other community members. The heterotrophs of the *Lokiarchaeia, Spirochaetes, Calditrichia* and *Bacteroidales* ferment these compounds producing acetate and hydrogen, which in turn can be used as energy source of ETH-SRB1 and Seep-SRB1c.

ANME-1-GMV encodes the reverse methanogenesis pathway (Table S7). As most ANME, ANME-1-GMV does not encode respiratory pathways that could provide sinks for the reducing equivalents released during methane oxidation. Instead, ANME-1 is expected to transfer these reducing equivalents to partner bacteria performing sulfate reduction(7,8). Strikingly, ANME-1-GMV encodes two alpha subunits of the methyl-coenzyme M reductase (mcrA), the key enzyme of the reverse methanogenesis. Both *mcrA* genes are highly similar and are located on the same scaffold, separated by 13000 bp (Figure 4B). One *mcrA* copy forms an operon with *mcrBG*, whereas the second *mcrA* copy occurs isolated. To our knowledge, this is the first report of an ANME-1 genome with two *mcrA* genes. Interestingly, the isolated *mcrA* is truncated (1.388 bp vs 1.691 bp) and flanked by two genes encoding tyrosine-type integrases and two genes encoding uncharacterized proteins with the domain COG4090. More specifically, the tyrosine-type integrase gene resembles a XerD-like integrase. XerC and XerD are tyrosine recombinases commonly found in bacteria, where they maintain chromosome topology(102). However, tyrosine-type integrases can also occur as proviruses within archaeal genomes(103), and there are reports of phages using the XerC/XerD machinery to integrate within the host genomes(104). Previous work on the ANME-1 virome provided evidence for virus-mediated gene displacement, including for the *thyX* gene(5,54). In the same study, *XerC/XerD* genes were present in several ANME-1 associated viruses(54). Moreover, recent studies highlight the complex phylogeny of MCR, with multiple lateral gene transfer events(105,106), including the proposal that ANME-1 acquired their *mcr* genes from the *Methanofastidiosa*(5). Taken together, we hypothesize that the observed mcrA duplication in ANME-1-GMV was mediated by a mobile genetic element or virus, although the duplicated *mcrA* appears truncated. Whether this truncation affects the protein function remains to be determined.

Notably, several other ANME-1 MAGs also contain truncated *mcrA* genes flanked by *XerC/XerD* (Figure 4B, Table S8). Initially, these ANME-1 genomes only seem to have the truncated *mcrA* copy flanked by *XerC/XerD*, while the *mcrBG* genes were usually located separately at the end of other contigs. However, upon closer inspection we detected partial *mcrA* fragments next to the *mcrBG* genes in the corresponding genomes. These gene fragments were not automatically annotated due to insufficient completeness and position at contig termini (Table S9). Actually, the genome GCA_030602585.1 (JAFNKG01 sp. 030602585, belonging to the same species as ANME-1-GMV) also harbors two *mcrA* copies forming the same operon configuration observed in ANME-1-GMV (Figure 4B). A phylogenetic analysis of the different mcrA revealed that ANME-1-GMV and ANME-1 JAFNKG01 sp. 030602585 mcrA sequences cluster together, indicating minimal evolutionary divergence between the complete and the truncated *mcrA* genes (Figure 4B). However, these two sequences do not form a monophyletic group with the remaining truncated *mcrA* genes identified in other ANME-1. Future studies should investigate the evolutionary history of these truncated *mcrA* genes, including their potential links to environmental pressures or virus-mediated gene transfer events.

### Metabolic capabilities of the putative partner SRB

The Seep-SRB1c-GMV genome is the only MAG in the biofilm that encodes a complete sulfate reduction pathway (Table S7), and therefore, likely represents the syntrophic partner of ANME-GMV1 in AOM. Seep-SRB1c-GMV is the second-most abundant organism in the biofilm (Figure 2B), its abundance is only one sixth that of ANME-1-GMV. However, reported similar differences in partner abundances have been observed previously in AOM consortia at genomic level(7). Interestingly, Seep-SRB1c bacteria have never been identified microscopically or in enrichment cultures as AOM partner bacteria. Still, a role of Seep-SRB1c as AOM partner of ANME-1 and ANME-3 has been hypothesized based on genomic co-occurrence(107). Seep-SRB1c-GMV encodes several multiheme cytochromes that could facilitate direct interspecies electron transfer (Table S7)(5,6). Notably, Seep-SRB1c-GMV also encodes a hydrogenase (Table S7; Seep-SRB1_00190-191), an unusual feature for most AOM partner bacteria(6,7). To date, only the thermophilic AOM partner bacterium “*Candidatus* Desulfofervidus auxilii” contains hydrogenases(15). Similarly, the hydrogenase of Seep-SRB1c-GMV might allow to couple sulfate reduction with hydrogen consumption. Such metabolic flexibility may enable Seep-SRB1c-GMV to survive in the absence of electron-donating ANME partners.

### The minimal side community has a heterotrophic lifestyle

Previous studies revealed the dominance of ANME-1 and partners in methane-oxidizing biofilms, but pointed out the presence of other microorganisms(21,24). In the GMV biofilm, the associated community constitute only ∼10% of the metagenomic reads (Figure 2B, Table 1). This side community is formed by five apparently clonal strains, which represents an ultralow biodiversity, and by this it is far less diverse than AOM enrichment cultures maintained in the laboratories for >10 years(108). However, biofilms and enrichment cultures share some clades like *Spirochaeta* and *Bacteroidetes.* Besides, some of these MAGs showed high identities to deep-sea marine organisms such as the Spirochaetia*-*GMV affiliated to *Sediminispirochaetaceae*(109–111), and the Loki-GMV MAG, closely related to *Prometeoarchaeum syntrophicum,* the first cultured asgardarchaeum, enriched from AOM cold seep sediments. Altogether, the side community encode different proteases and carbohydrate-active enzymes (Supplementary Information, Tables S10 and S11) as well as hydrogenases (Table S7) that suggest a potential to ferment organic compounds or by-products released by the AOM partners, as indicated by the specific ranges in δ^13^C values of the bacterial lipids (Table 2). Their dependence on the ANME and Seep-SRB1c organisms may explain their low genomics abundances.

### 7 A fluid path sustains a minimal AOM biofilm community

Our analyses revealed that a subsurface biofilm had formed at an interface, where sulfate intrudes from above and below. The biofilm consists of a remarkably simplified microbial community. Based on genomic and lipid evidence, ANME-1b mediates methane oxidation, which appears to be coupled to sulfate reduction performed by Seep-SRB1c-GMV. The presence of Seep-SRB1c as a potential partner bacterium is noteworthy, as it strengthens previous evidence that this clade represents a novel group of AOM partner organisms(107). Targeted cultivation and microscopy studies are required to confirm the proposed interaction of these organisms.

Typically, AOM occurs, where substantial sulfate and methane concentrations overlap, and steep gradients supply large amounts of substrates to the microorganisms. In surface sediments this may result in cell densities ranging from 10^6^ to 10^10^ ANME cells cm^-3^ depending on the environment(17). In contrast, an active AOM biofilm appears at a depth, where sulfate is largely depleted (Figure 1). Diffusion alone would generate very low sulfate and methane fluxes (∼0.1 mmol m² day^-1^) into about 10-meter-deep AOM zone, which could support only a very limited number of microbial cells. However, in this location, sulfate penetrates from above and below. The accumulation of large visible biomass suggests that advective transport dominates. Indeed, the biofilm locates at the rim of the GMV. Here, high fluid pressures in the mud volcano feeder channel may have resulted in cracks which favor and focus methane migration in the surrounding hemipelagic sediments, as previously suggested based on geochemical and geophysical observations(44). Notably, methanotrophic biofilms occurring in sulfate-depleted horizons have been previously reported as a result of steady supply of methane and sulfate(23).

Only a few examples of AOM-fueled biofilms have been described(18–24), and those occurring within fractured sediments are usually dominated by ANME-1b(21,22,24). Similarly, our biofilm community is overwhelmingly dominated by ANME-1b, while the surrounding sediment showed lower abundances of this clade (Figure 2). Interestingly, a previous AOM enrichment from GMV sediment using a biotrickling filter yielded a community dominated by of ANME-1b, despite this clade being rare in the inoculated sediment sample(112). The recurrent finding of ANME-1b-dominated biofilms suggests that at least some members of this clade might possess the ability to form microbial mats. Actually, we could identify a distinct archaeal lipid composition, particularly a high abundance of acyclic and macrocyclic PE-archaeols, that might be related to the formation of such thick biofilms. We searched in the ANME-1-GMV genome for specific adaptions in the lipid biosynthesis pathway (Supplementary Information, Table S7, Figure S1). ANME-1-GMV encodes the canonical archaeal route for lipid biosynthesis, although gene duplications in certain steps may indicate functional diversification, as previously proposed(113). For instance, ANME-1-GMV possesses several genes for ring synthase (GrsA/GrsB; Supplementary Information, Figure S2), which might explain the proportionally high abundance of macrocyclic PE archaeols next to the more common PG-GDGTs (Figure 3, Table S4).

The side community of the biofilm consist of heterotrophs in low abundance likely feeding on biomass or exudates released by the AOM core community (ANME and SRB). Previous studies on methane-oxidizing biofilms and enrichments have shown that the AOM partners can sustain other microorganisms(99,108). Interestingly, several clades are shared across samples, suggesting the existence of a core AOM-associated side community that includes groups like *Spirochaetia*, *Bacteroidales* and *Lokiarchaeota*, the latter of which has been repeatedly found in SMTZs(82,107,114,115). The potential metabolic interdependencies among the members of the biofilms could require specific spatial and environmental arrangements. Such conditions might promote potential events of horizontal gene transfer (HGT). Therefore, biofilms have been proposed to be hotspots for HGT events(116,117). In this context, the presence of two *mcrA* genes in the ANME-1-GMV MAG flanked by putative integrases could indicate a past viruses-mediated HGT event. These events could also occur among minor components of the microbial community. Indeed, growing evidence indicates that numerous HGT events have shaped the evolution and diversification of Asgardarchaeota(118,119). Future studies should investigate the role of HGT among microorganisms inhabiting sulfate-methane transition zones.

## Conclusions

In deep anoxic sediments at GMV, we described a localized SMTZ harboring a microbial biofilm. It has an ultra-low diversity microbial community with only seven different microorganisms, but strongly dominated by an archaeal species of ANME-1b. This ANME is most likely syntrophically associated with a SeepSRB-1c. Other lower-abundance microorganisms have the genomic potential to use different organic products, probably forming a metabolically interdependent network both among themselves and with the AOM partners. Lipidomic analysis revealed the presence of distinct macrocyclic archaeols in this biofilm, probably synthesized by the ANME-1b, what might indicate a potential adaptation of this clade to form biofilms, since this group has been repeatedly found in other AOM biofilms. The presence of fluids pathways at the rim of the GMV adds evidence to an extensive fracture network around MVs edifices, which channel substantial amounts of methane-rich fluids through the subsurface. At mixing zones with sulfate-bearing waters, dense AOM microbial communities establish, which mitigate the methane flux to the surface. These new findings challenge the common belief that MV summits are the most microbially active areas and prompt a reevaluation of the outer limits of such features in the search for signs of microbial life.

## Acknowledgements

We want to thank captain, crew, and scientists of expedition M149 for the help onboard and for the excellent data produced during the cruise. We also thank Kai Hachmann for extraction and preparation of the samples for lipid analyses and Ana Gutiérrez-Preciado for her help with the metabolic analysis of the side community.

## Data availability

Generated sequences and MAGs were deposited under NCBI BioProject ID PRJNA1391031. During the revision process, the read libraries can be consulted under the following reviewer <u>link</u>. Currently, MAGs and additional data such as phylogenetic trees and alignments are available at the following link: https://saco.csic.es/s/BZ4XCniZSadLBoD

