## Supplementary Text for "Anaerobic oxidation of methane supports a minimal microbial community in a Subsurface Biofilm at Ginsburg Mud Volcano"

**Supplementary Information**

**Supplementary Methods**

Analysis of the 16S rRNA gene libraries.

Archaeal and bacterial 16S rRNA gene libraries were analyzed according to Laso-Pérez *et al.* (2019)^1^. In short, primer and adapter sequences were removed using Cutadapt (v.1.9.1)^2^ with no indels and 0.16 as the maximum allowed error rate (-e). Archaeal reads were first merged with PEAR^3^ (v.0.9.6; 10 bp as minimum overlap and 400 and 570 bp as minimum and maximum assembly lengths) and them trimmed with Trimmomatic (v0.35)^4^ with 6:12 as sliding window and 450 bp as minimum length. Bacterial reads were first trimmed (sliding window 4:15; and 100 bp as minimum length) and then merged (minimum overlap 10 bp; maximum length 500 bp). Processed reads for bacteria and archaea were dereplicated and clustered into operational taxonomic units (OTUs) using Swarm (v2.2.2; parameters: “*-b 3, -d 1, -f”*)^5^. Afterwards, OTUs were classified using SINA aligner (v1.2.11)^6^ against the SSURef_NR99_123 SILVA database and analyzed using the R software.

**Supplementary Results**

Metabolic potential of the side community

In the GMV biofilm, the associated community constitute only ~10% of the metagenomic reads (Figure 2B, Table 1) and is formed by five apparently clonal strains, some of which are related to deep-sea clades and shared with enrichment cultures. Spirochaetia*-*GMV is closely affiliated to *Sediminispirochaetaceae,* which is an anaerobic marine bacterium capable of using different organic compounds^7–9^. Similarly, the Bacteroidales*-*GMV encode a high number of proteases and carbohydrate-active enzymes (CAZymes), only comparable to the Calditrichia-GMV (Tables S10 and S11). Hence, it is likely that both organisms grow as heterotrophs in the biofilm, likely on biomass or cell exudates produced by the AOM core community, as also indicated by the specific ranges in δ^13^C values of the bacterial lipids (Table 2), and previously shown for AOM enrichments^10^. These three organisms encode hydrogenases. The Spirochaetia-GMV possess an operon for a FeFe-hydrogenase Hnd, while the *Calditrichiota* and the *Bacteroidales* genomes encode NiFe hydrogenases (Table S7). As fermenting microorganisms, these hydrogenases would likely produce hydrogen. Alternative, some of these organisms could produce acetate as fermentation product (Table S7).

The ETH-SRB1-GMV genome contains less genes encoding proteases and CAZymes compared to the other bacterial genomes*.* Interestingly, ETH-SRB1-GMV does not encode a sulfate reduction pathway, which is present in closely related organisms^11,12^. This absence might be related to the completeness of the MAG (70%), which was the lowest of the whole dataset. Instead, we found a gene for a putative sulfide:quinone oxidoreductase (SQR), which is used for sulfide oxidation. The putative function of this enzyme under anoxic conditions is unclear.

The Loki-GMV MAG is closely related to *Prometeoarchaeum syntrophicum* (class *Lokiarchaeia*), and encodes a similar metabolic potential (Table S7). *P. syntrophicum* represents the first cultured asgardarchaeum. It was enriched from cold seep sediments using a continuous bioreactor, which favored the formation of AOM biofilms. This shows that members of the *Lokiarchaeia* are capable of growing as side-community organisms in AOM-dominated environments. The isolation of *P. syntrophicum* in amino acid-rich medium revealed that it functions as protein-degrading fermenter. As fermentation products formate and hydrogen are released into the medium and can serve as substrate of sulfate-reducer or methanogens^13,13^. The Ginsburg biofilm does not contain methanogenic archaea. Therefore, we hypothesize that Loki-GMV produces hydrogen, which may be used as additional energy source by Seep-SRB1c-GMV, the only MAG in the biofilm encoding a sulfate reduction pathway. Altogether, the MAGs of the side community have the potential to degrade organic compounds or by-products released by the AOM partners (Figure 5). Their dependence on the ANME and Seep-SRB1c organisms may explain their low genomics abundances.

Four MAGs of the biofilm (ETH-SRB1-GMV, Calditrichia-GMV*,* Bacteroidales-GMV and Seep-SRB1c-GMV) encode hypothetical pathways to reduce diverse nitrogen species (Table S7). This is notable, because they inhabit sediment horizons where oxidized nitrogen compounds should be depleted. For instance, Calditrichia-GMV has an operon encoding a dissimilatory nitrate reduction pathway that has been reported in another member of the clade, *Caldithrix abyssi*^14^. This operon encodes for a nitrate reductase of the Nap family, putatively responsible for the reduction of nitrate to nitrite, and for a periplasmic c-type octaheme oxidoreductase 𝜀Hao enzyme. This enzyme traditionally catalyzes the reduction of hydroxylamine to nitrite, but in *C. abyssi* 𝜀Hao was proposed to mediate the conversion of nitrite to ammonia^15^, maybe coupled to hydrogen oxidation. By contrast, the Bacteroidales-GMV genome encodes an Nrf complex, complex catalyzing nitrite reduction to ammonia. Furthermore, the ETH-SRB1-GMV also encodes another Nap operon encoding a putative nitrate reduction complex; and a NosZ allowing the reduction of nitrous oxide to nitrogen. Additionally, both, ETH-SRB1-GMV and Seep-SRB1c-GMV, genomes contain operon encoding two cytochromes resembling a NapC of nitrate or TMAO reductase and a hydroxylamine dehydrogenase. However, their functions in these organisms are unclear. Potentially they may form a nrfAH-like complex. The diversity of enzymes involved in the nitrogen cycle in our dataset is striking, because the studied subsurface sediments should be depleted in oxidized nitrogen species. Some of these enzymes might be catalyzing a different process. However, for some of them the whole operon structure is similar to other organisms where nitrogen cycling has been shown (i.e. Nap operon in the Calditrichia-GMV)^15^. Still, the genetic machinery might be present, but not transcribed. Another option is that these enzymes catalyze reactions involving other compounds, but resembling nitrogen-cycling enzymes. Finally, the encoded enzymes might be functional and reflect the potential of these organisms to live at different sediment horizons, where alternative electron sinks are available.

Lipid synthesis pathway in archaeal MAGs

To explain the relationship between some of these lipids with archaea in the biofilm, we searched for candidate genes encoding the potential lipid biosynthesis pathways. As most ANME-1 genomes, the ANME-1-GMV MAG encodes the complete pathway for archaeal lipid biosynthesis, including the steps to synthesize archaeol, and glycerol dibiphytanyl glycerol tetraethers (GDGTs) (Table S7, Figure S1). We also found potential candidate genes for the tetraether synthase (Tes), which has been recently identified as a critical enzyme for the production of GDGT from archaeol, but it also produces macrocyclic archaeol as a by-product^16,17^. Actually, many ANME-1 genome including ANME-1-GMV contain several gene copies encoding specific catalytic steps GDGT synthesis, what might be related to functional diversification as suggested in previous studies^18^. As examples, several genomes encode diglyceryl glycerol phosphate synthase (DGGGP). We also detected a putative gene encoding a GDGT cyclization protein or ring synthase (GrsA/GrsB; Figure S1, Table S7). Ring synthases are responsible to introduce specifically rings at different positions of the GDGT molecule in the archaeon *Sulfolobus acidocaldarius*^19^. A phylogenetic analysis of the *grs-*like proteins in archaea that included the two sequences from archaeal species in the biofilm showed that the potential homolog in *Lokiarchaeia* species was located in a different clade from any grs-like sequences, while the homologue of ANME-1-GMV MAG clustered them with other putative ANME-1 *grs,* but in a sister clade of canonical *grs* of *S. acidocaldarius*. Therefore, it is unclear if the candidate *grs* of ANME-1 function as canonical enzymes. Alternatively, they might be involved in other cyclization reactions. For instance, our lipid analysis detected macrocyclic archaeols with rings, which might have been produced by the corresponding putative Grs-like proteins. Our genomic analysis could find all genes necessary for lipid biosynthesis, but we were not able to directly link any ANME-1-GMV gene to the distinctive lipid composition of our samples. Still, the duplication of several enzymes of the lipid biosynthesis pathway and the widespread presence of *grs* homologues suggests the ability of ANME-1 to produce different lipids that might contribute to their adaptation through changes in membrane rigidity and thermal stability to different environments, temperatures or lifestyles, i.e. forming large biofilms.

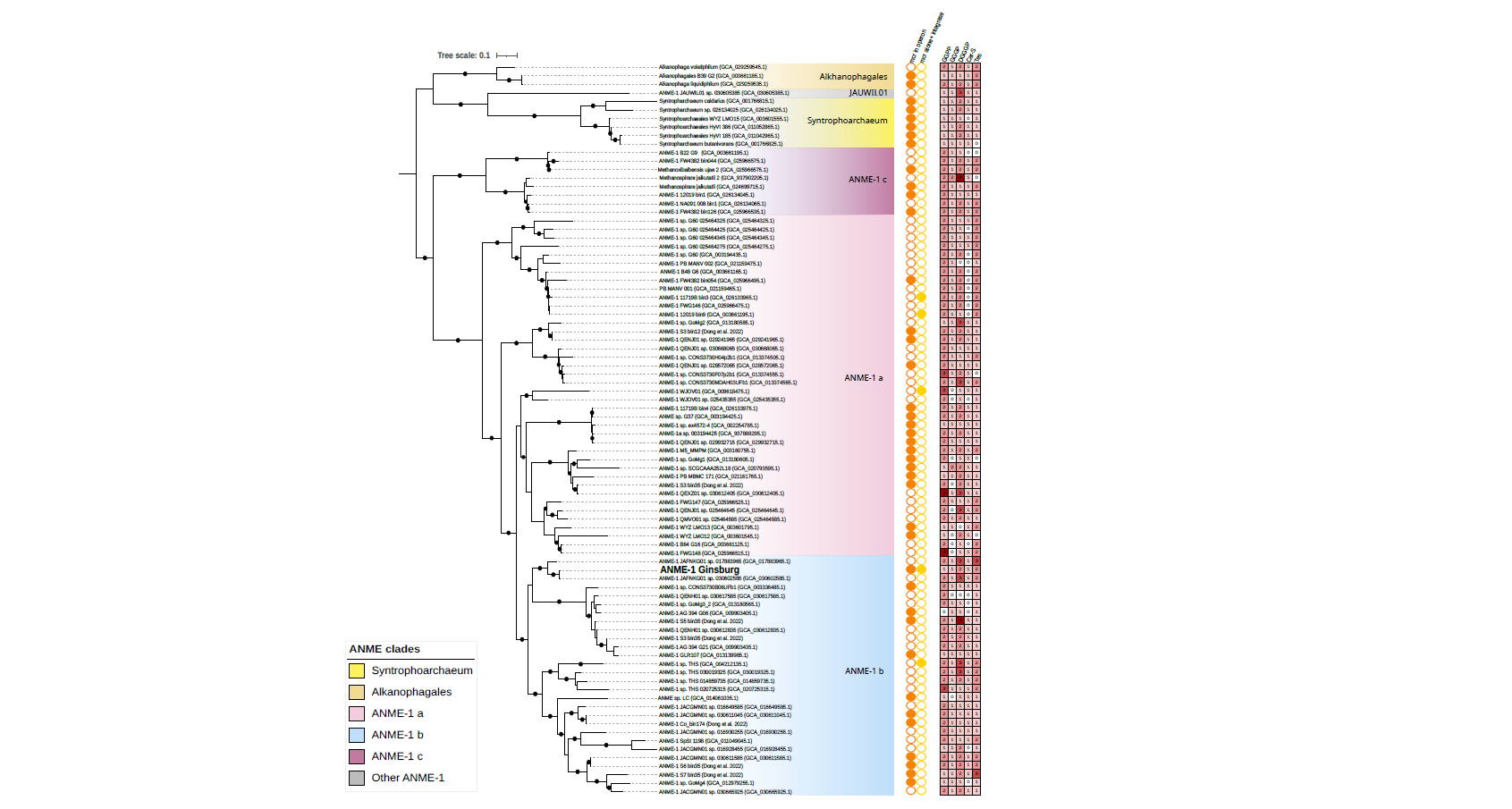

**Figure S1.** Maximum-likelihood phylogenomic tree based on 105 MAGs from the class Syntrophoarchaeia, showing the presence/absence *mcr* genes, archaeal lipid biosynthesis genes and synteny analysis of genomic regions harboring mcr operons across representative MAGs. ANME of interest clustering closely with ANME-1b and JAFNKG01 sp. 030602585. GGPP: geranylgeranyl diphosphate synthase; GGGP: geranylgeranylglyceryl phosphate synthase; DGGGP: digeranylgeranylglyceryl phosphate synthase; car-S: carboxy-SAM synthase; Tes: tetraether synthase.

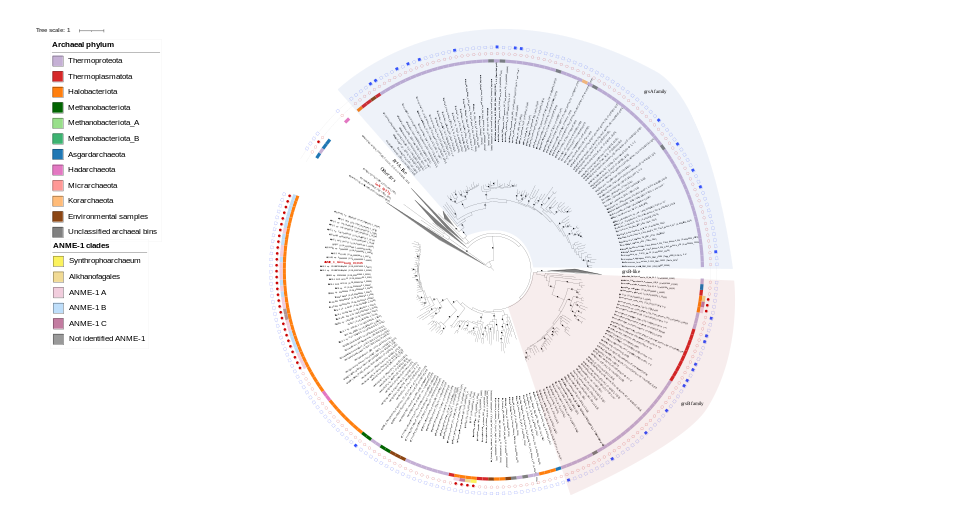

**Figure S2.** Phylogenetic tree of grsA and grsB homologs in archaea. Red circles indicate genomes newly added in this study that encode grs homologs. Blue squares represent isolated archaeal strains that have been shown to produce GDGTs, even though a specific grs-like enzyme has not been isolated. Taxonomic classification based on GTDB, along with the historical ANME-1 clade designation is shown in the color strips. Asterisks (*) denote the specific GDGTs produced by each organism. GrsA and grsB sequences characterized in the original study are shown in bold and the two archaeal sequences from the biofilm species in bold-red. Members of the phylum Halobacteriota (mostly ANME-1a and ANME-1b groups) form a distinct monophyletic clade whose grs-like homologs do not cluster with canonical grsA or grsB-like proteins from other archaeal phyla, suggesting the presence of a potentially novel clade of grs homologs.
