## Supplementary material for "Anaerobic oxidation of methane supports a minimal microbial community in a Subsurface Biofilm at Ginsburg Mud Volcano": Figure S1

**Figure S1**. Maximum-likelihood phylogenomic tree based on 105 MAGs from the class *Syntrophoarchaeia*, showing the presence/absence mcr genes, archaeal lipid biosynthesis genes and synteny analysis of genomic regions harboring mcr operons across representative MAGs. ANME of interest clustering closely with ANME-1b and JAFNKG01 sp. 030602585. GGPP: geranylgeranyl diphosphate synthase; GGGP: geranylgeranylglyceryl phosphate synthase; DGGGP: digeranylgeranylglyceryl phosphate synthase; car-S: carboxy-SAM synthase; Tes: tetraether synthase.

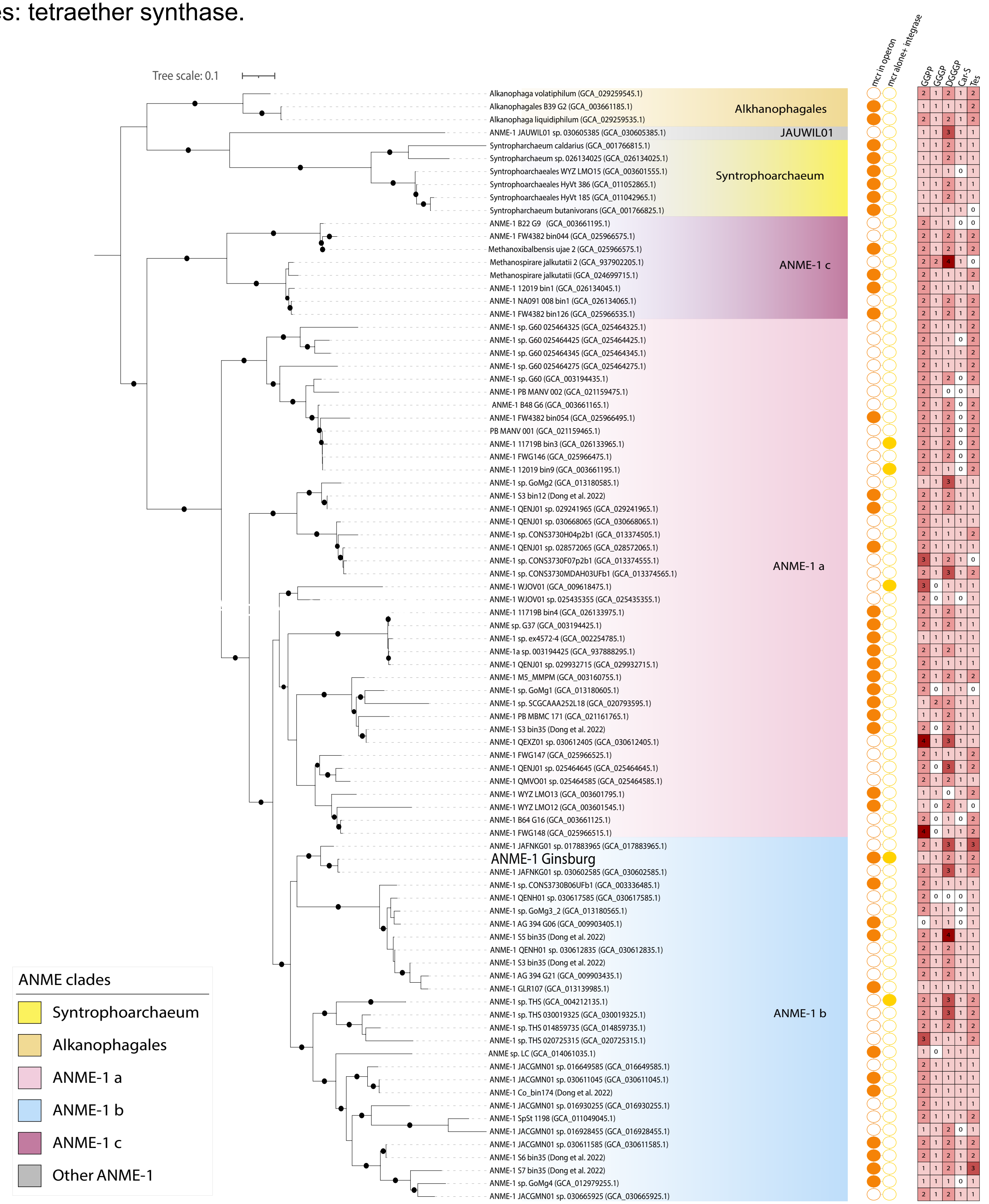
