## Supplementary material for "Anaerobic oxidation of methane supports a minimal microbial community in a Subsurface Biofilm at Ginsburg Mud Volcano": Figure S2

**Figure S2**. Phylogenetic tree of grsA and grsB homologs in archaea. Red circles indicate genomes newly added in this study that encode grs homologs. Blue squares represent isolated archaeal strains that have been shown to produce GDGTs, even though a specific grs-like enzyme has not been isolated. Taxonomic classification based on GTDB, along with the historical ANME-1 clade designation is shown in the color strips. Asterisks (*) denote the specific GDGTs produced by each organism. GrsA and grsB sequences characterized in the original study are shown in bold and the two archaeal sequences from the biofilm species in bold-red. Members of the phylum *Halobacteriota* (mostly ANME-1a and ANME-1b groups) form a distinct monophyletic clade whose grs-like homologs do not cluster with canonical grsA or grsB-like proteins from other archaeal phyla, suggesting the presence of a potentially novel clade of grs homologs.

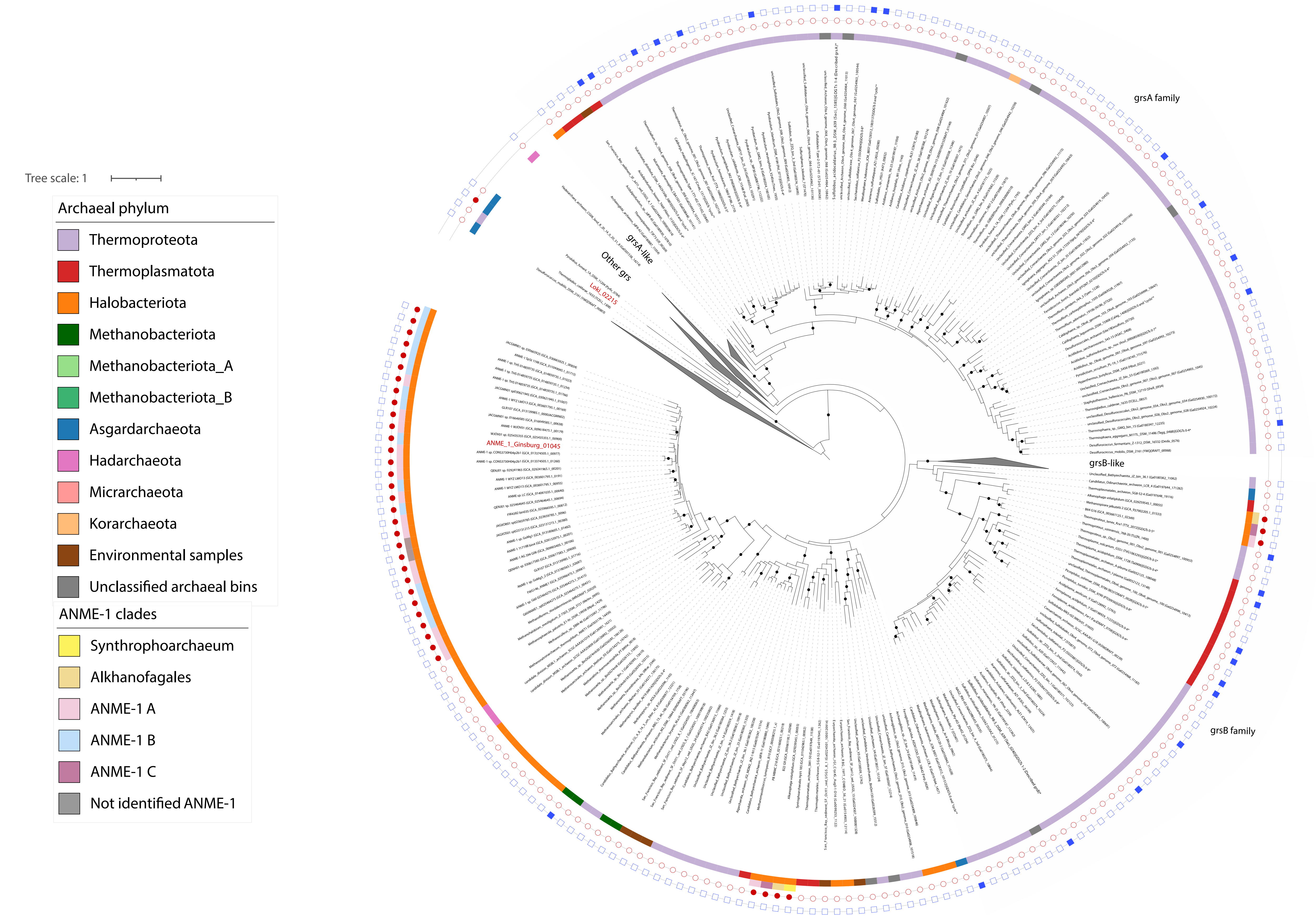
